# Phyllotaxis Models: from the Inhibition Potential to the Real Plant

**DOI:** 10.1101/2021.08.08.455557

**Authors:** Walch Jean-Paul

## Abstract

Previous phyllotaxis models allowed the initiation of new primordia when a threshold of inhibition potential is reached on the meristem front: their adequacy to botanical reality is only qualitative. We formulated the hypothesis that it is not the value of the inhibition threshold that remains constant as the meristem develops, but the difference of the inhibition thresholds during the initiation of two successive primordia. We were thus able to model with accuracy the sequence of plastochron ratios observed by Williams (1975) on the leaf meristem of flax: an outstanding result. More generally, we have shown that the evolution trajectories of the phyllotaxis modes as a function of the plastochron ratios follow the minima of the potential under decreasing plastochron ratios constraint and bifurcate when the number of these minima increases, thus giving physicochemical foundations to the famous van Iterson diagram. This historical representation of rising phyllotaxis shows the trajectories, but doesn’t give the velocity of the movement: our plastochron ratio sequence adds this major dynamical information.

## 1 Introduction

The superb spirals generated by plants have fascinated generations of botanists before mathematicians succeeded in modeling them (Mitchison, 1977; Douady and Couder, 1992, 1996; Smith and al., 2006; Atela, 2011; Golé and al., 2016; Douady and Golé, 2016). But, the adequacy of the theory with the observations of botanists has most often been only qualitative.

It was not until the end of the 20th century that Douady and Couder (1992, 1996) developed the standard model. The future organs, leaves, petals, stamens or carpels originate within the meristem, at the tip of the bud, as small primordia which are initiated in a crown or front some distance from the tip of the meristem (apex) before to migrate down its ogival surface. As early as the 19th century, Hofmeister (1868) had an intuition of the rule for the appearance of a new primordium: it is inserted as soon as the space left available by its predecessors located on the front is large enough. Standard theory models an inhibition field produced by young primordia that prevents the formation of a newcomer as long as the intensity of the inhibition potential is too high. The centrifugal movement of these potential sources due to the expansion of the meristem is responsible for a decrease in the field intensity on the front, so that a new primordium can be initiated when the potential at a point become below a certain threshold. It is at this point that the new primordium is initiated, and the succession of these produces the famous spirals without any other kind of intervention.

Most models (Douady and Couder, 1996b; Smith and al., 2006) thus consider that a new primordium is formed when the intensity of the inhibition field at the level of the front is below a threshold whose value does not change when the meristem grows. However, in botanical reality, during the development of the meristem, the primordia are more and more numerous and more and more crowded on the front (figure 1), which necessarily increases the inhibition potential. In one of their models, Douady and Couder (1996b) vary this threshold, and observe that the results are qualitatively similar to those obtained with a constant threshold: the different modes of spiral phyllotaxis are obtained as well as the evolution during the development of the meristem towards higher and higher modes corresponding to increasing numbers of spirals.

**Figure 1.**
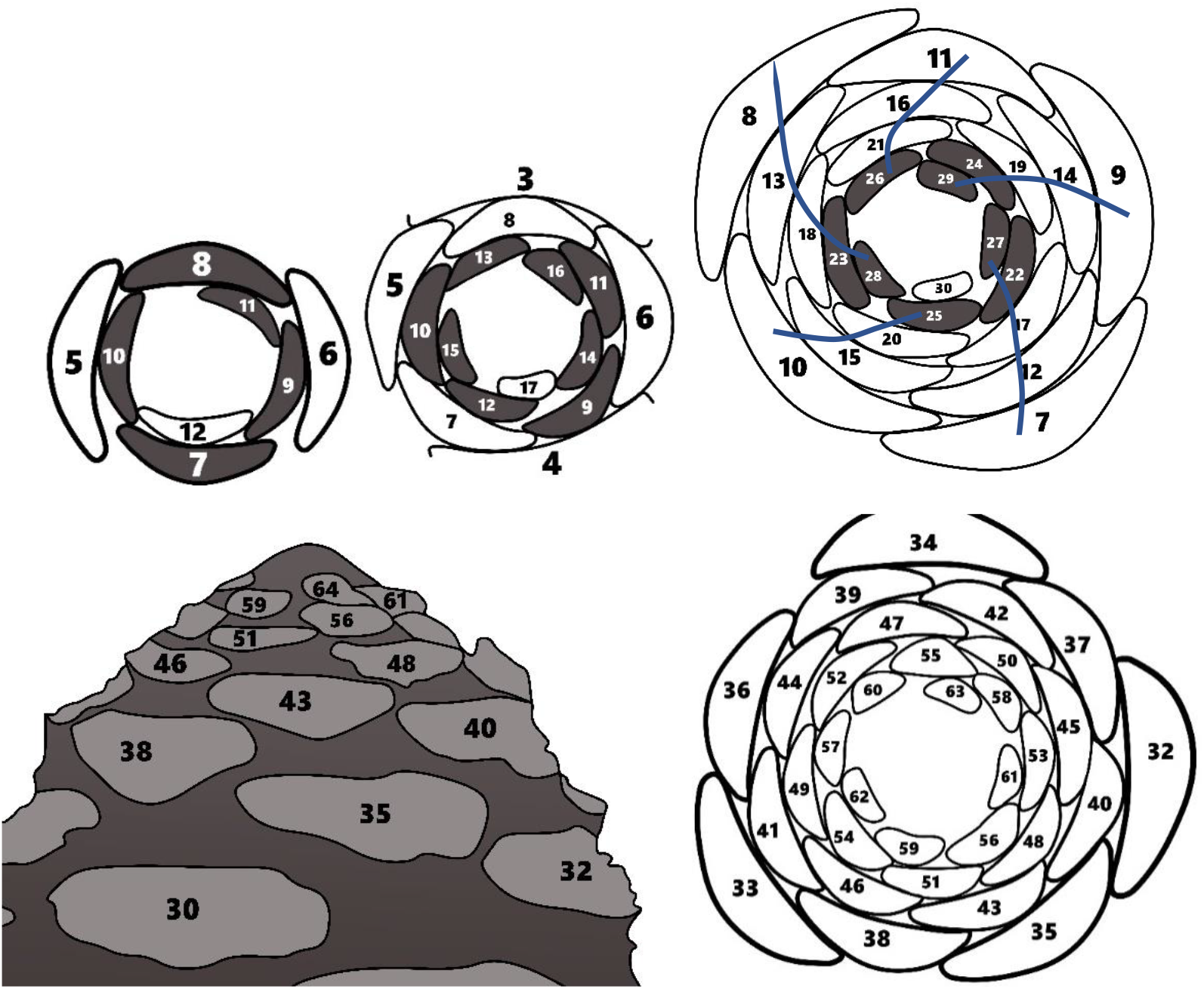
The meristem of *Linum usitatissimum* observed by Williams (1975) for the initiation of primordia 12, 17 and 30 between the 11th and 15th days of growth. The primordia (future leaves) are more and more tight on the front (in black): their number goes from 5 to 8. The phyllotaxis mode which corresponds to the number of pairs of dextral and sinistral spirals (parastichies) as well as to a pseudo-orthostichy, goes from (2,3.5) to (3,5,8) then (5,8,13). The primordia are numbered according to their chronological sequence of initiation. The divergence angle is the angle between two successive primordia. It is close to 180° for primordia 5 and 6, and to 137° for numbers 22 and 23. Linum usitatissimum meristem according to Meicenheimer (2006). Section of bud at the age of 22 days according to Williams.

The diameter of the meristem increases over time. If d_i_ is the distance of primordium i from the apex and d_i + 1_ that of its chronological successor (i + 1), the “plastochron ratio” is the ratio (Richards, 1951):

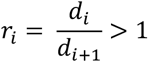

The increasingly dense concentration of primordia on the front corresponds to a regular decrease in the plastochron ratio during the growth of the meristem: each new primordium is found closer to its neighbor.

The hard disk model was used by van Iterson (1907) to formalize his famous diagram in phase space (divergence angle, plastochron ratio) which shows the rise of phyllotaxies when plastochron ratios decrease. The construction method consists of filling a cylinder with disks of equal diameters. We then observe parastichies conforming to Fibonacci spirals. The system goes up the Fibonacci sequence if the diameter of the discs decreases compared to that of the cylinder.

Godin, Golé and Douady (2020) interpret the development of a plant as a “ travel” down the van Iterson diagram. Monocots and dicots start on a branch of the diagram low mode, for instance (1,1), with the second primordium opposite the first. Then the plastochron ratio decreases, which leads to the choice of a pattern (1,2) or (2,1) corresponding to the chirality of the plant (dextral or sinistral). If the branch (2,1) is chosen, the “Fibonacci rule” imposes the phyllotaxis (2,3). The decrease in the plastochron ratio then inexorably leads to increasingly high Fibonacci modes.

Williams (1975) measured this evolution on the leaf meristem of *Linum usitatissimum*.

In this paper, we have accurately modeled the meristem of flax as drawn by Williams at the different ages of its growth, including the evolution of parastichies with the decrease in plastochron ratio. In addition, the divergence angle between two consecutive primordia rapidly converges to the golden angle (137.5°).

The classic hypothesis (Douady, 1996; Smith and al. 2006) according to which a new primordium is formed when the intensity of the inhibition field at the level of the front is below a threshold whose value does not change during growth of the meristem is not verified, but on the contrary, it is the difference in the inhibition thresholds during the initiation of two successive primordia which seems constant.

We limit our study to the case where the range of the inhibition field is short, which corresponds to primordia behaving almost like hard disks whose inhibition prevents interpenetration: a classic hypothesis. In this case, all the calculations can be derived analytically. We then demonstrate that a constant value of the difference in inhibition potential between two successive primordia generates Fibonacci phyllotaxis as well as the plastochron ratios observed by Williams on flax that we call “canonical sequence” of plastochron ratios.

To obtain this result, we had to calculate the inhibition potentials as a function of plastochron ratios and divergence angles. It appears that the evolution of phyllotaxis modes during the decrease of plastochron ratios climbs the potential “talwegs”. This result is consistent with the van Iterson diagram and gives it physicochemical foundations.

For lower and lower plastochron ratios, the potential function presents a growing number of minima, which generates bifurcations between phyllotaxis modes also predicted by the geometric model of van Iterson.

The paper is structured as follows. In section 2, we present the mathematical model. This allows us to model the *Linum usitatissimum* meristem in section 3. Calculation of potential values gives physicochemical foundations to the van Iterson diagram (section 4). In section 5, the inhibition threshold difference hypothesis allows us to reproduce the *Linum usitatissimum* platochron ratio sequence. This opens up new insights for phyllotaxis research.

## 2 The model

Following the standard model (Douady and Couder, 1996; Smith and al., 2006; Kitazawa and Fujimoto, 2015) we describe the initiation of primordia as depending of the intensity of an inhibition potential on the meristem front.

The meristem front is represented as a circle with radius R_0_, the primordia being points located between this front and the outer edge of the meristem. The initiation of a new primordium takes place on the front at the point of polar coordinates (R_0_, θ), the initiation angle being that which minimizes an inhibition potential U_i_ generated by the already existing primordia. This potential decreases exponentially with the distance from the generating primordium, the decay length being γ. On the other hand, the potential generated by each primordium can decrease exponentially with the age of the generating primordium, the decay rate α_i_ being able to be different from one primordium to the other.

If d_ij_ is the distance between the new primordium i and the already existing primordia j (between 1 and i-1) located at the distance r_j_ from the center and with angular coordinate θ_j_ :

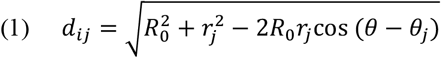

the potential is (2):

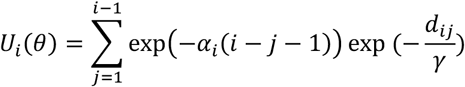

**Figure 2.**
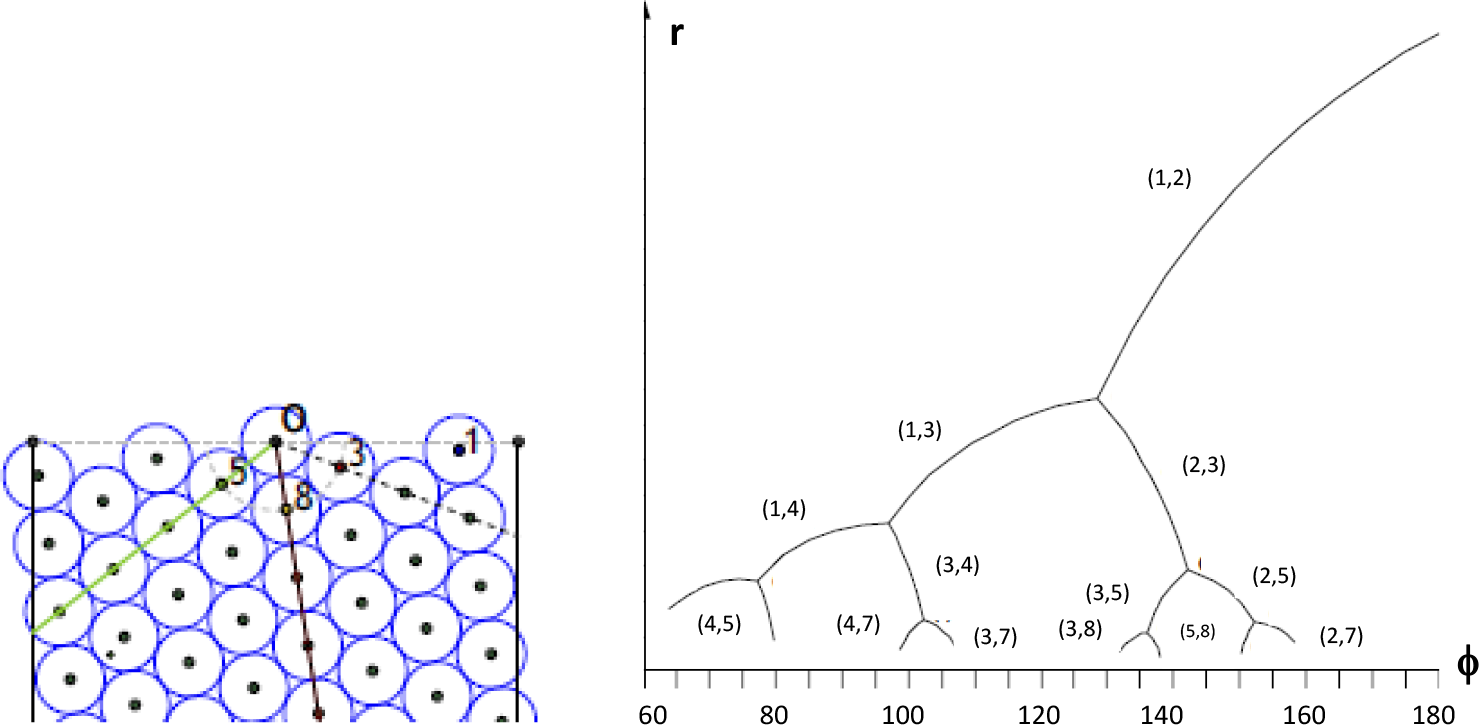
The stacking of hard disks in a cylinder produces a spiral phyllotaxic pattern of mode (3,5,8) for a certain ratio of the diameter of the disks to that of the cylinder (Godin and al., 2020); the van Iterson diagram represents the trajectories of phyllotaxis modes as a function of the divergence angle (ϕ) in degrees when the plastochron ratio (r) decreases.

For each species, the model parameters are the plastochron ratios between each pimordium i and its immediate successor and the α_i_ coefficients.

The angular distance θ – θ_j_ is a multiple (j-i) of the divergence angle if this one is constant. On the other hand, if the plastochron ratio is constant, the distance to the center of the primordium j is equal to r_j_^j^. Under these conditions we can calculate by equation (1) the distance d_ij_ of all the primordia already existing to the new primordium.

For a divergence angle of 137.5° (Fibonacci angle), the primordia whose angular distance to the new primordium modulo 360° is the smallest are, in decreasing order of the absolute value of the angular distance, the numbers (i + 3), (i + 5), (i + 8), (i + 13),… The terms of the Fibonacci sequence are recognized. According to equation (1), with a positive cos (θ – θ_j_), the distance d_ij_ of the old primordia increases with the plastochron ratio r_j_ and decreases with the angular difference (θ – θ_j_).

Therefore, for high plastochron ratios (eg: 1.3), the primordium (i + 3) will be closest to the new primordium, then it will be the primordium (i + 2): the phyllotaxis mode will therefore be (2,3). When the plastochron ratio decreases, it will be stronger angular differences which will minimize the distances. For r = 1.04, these will promote the primordia (i + 8) and (i + 5): the phyllotaxis mode will be (5.8).

**Figure 3.**
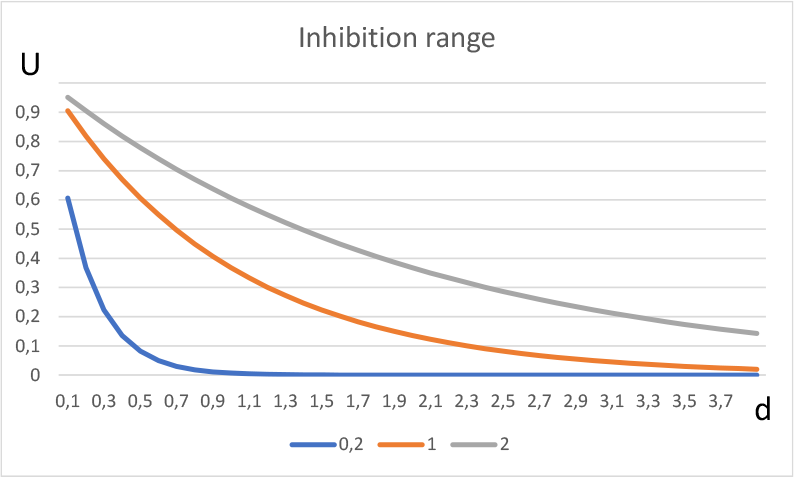
Range of the inhibition field according to the decay length: γ.

The coefficient γ, or decay length, determines the range of the inhibition. For γ = 0.2, the primordia behave almost like hard disks and the new primordium simply fits into the larger physical space left available by its predecessors.

If γ is high, the primordia produce a long-range field, with all existing primordia participating in the inhibitory potential at any point on the meristem.

## 3 Modeling of Linum usitatissimum *meristem*

We use data from Williams (table 1) to model the *Linum usitatissimum* meristem, and complete them with values of decrease in inhibition potential with age for both cotyledons (α_1_=α_2_=1), which is necessary to form a spiral phyllotaxis. The decay length is set to γ = 0.2.

**Figure 4.**
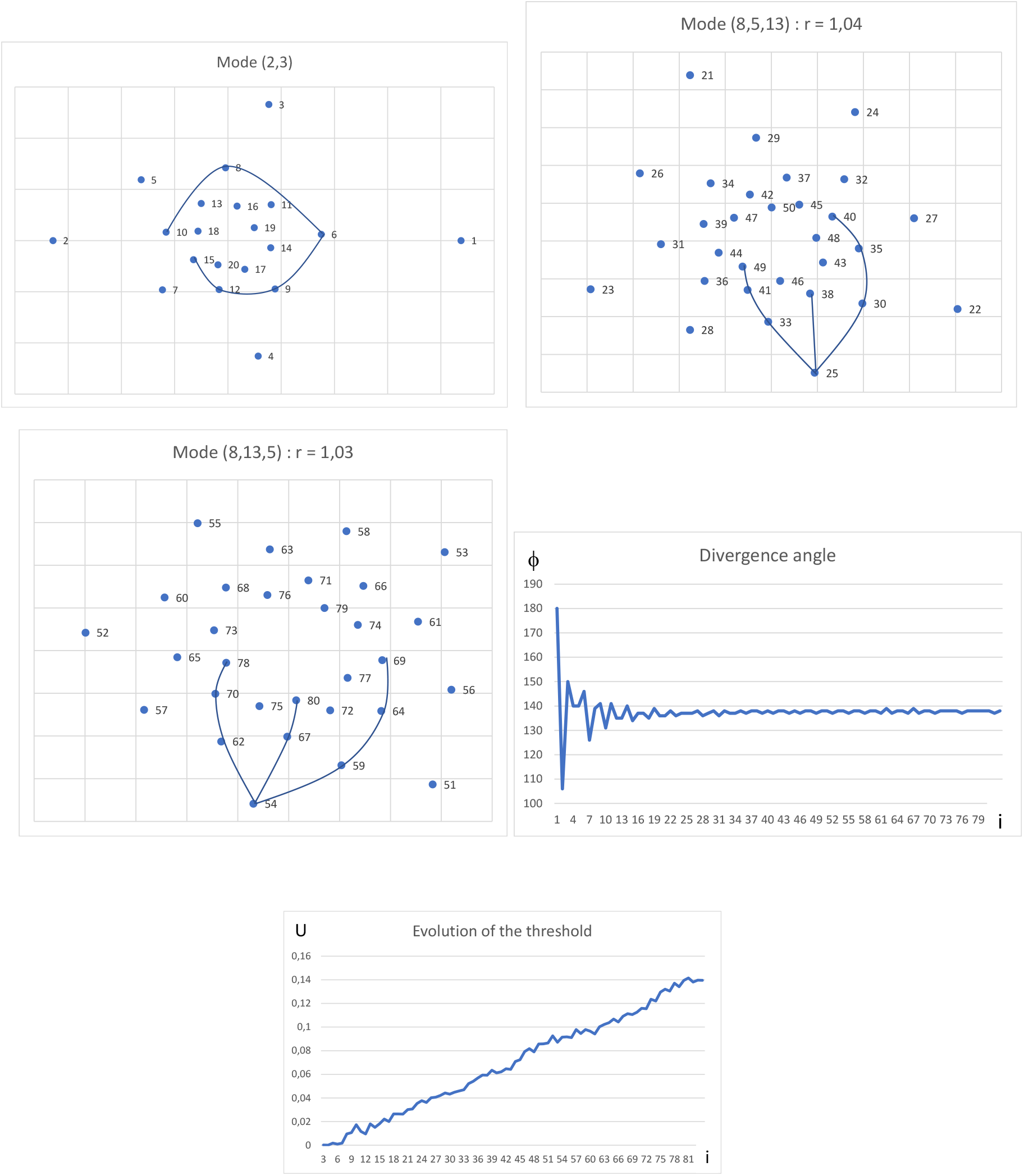
Modeling of the *Linum usitatissimum* meristem (compare to figure 1); divergence angle and inhibition threshold according to the primordium number.

**Table 1.** Evolution of the plastochron ratio with the age of *Linum usitatissimum* plant according to Williams (1975).

| Age (days) | Primordium interval | Plastochron ratio |  | Age (days) | Primordium interval | Plastochron ratio |
| --- | --- | --- | --- | --- | --- | --- |
| 1 | Cot.-2 | 1.330 |  | 25 | 63-73 | 1.045 |
| 4 | 2-4 | 1.189 |  | 29 | 84-94 | 1.041 |
| 4 | 4-6 | 1.152 |  | 32 | 100-110 | 1.044 |
| 8 | 4-14 | 1.101 |  | 36 | 125-135 | 1.035 |
| 11 | 10-20 | 1.082 |  | 39 | 151-161 | 1.030 |
| 15 | 20-30 | 1.073 |  | 43 | 173-183 | 1.033 |
| 18 | 35-45 | 1.052 |  | 50 | 202-212 | 1.034 |
| 22 | 51-61 | 1.049 |  |  |  |  |

The model accurately reproduces the flax meristem disposition, and the divergence angle rapidly converges to the golden angle. Remarkably, the inhibition threshold is also rapidly a linear function of the primordium number (slope = 0.00178, R^2^ = 0.994).

## 4 Potential values

We use an algorithm which allows to define a diagram which we call “phyllotaxis frieze” of phyllotaxis regions in phase space (ϕ, r). For a given plastochron ratio and divergence angle we calculate the distances from all primordia to the first of them (as in Table 2). We retain the two closest primordia: thus, for a divergence angle of 137.5° and a plastochron ratio of 1.04, the phyllotaxis will be in a mode (5.8).

**Figure 5.**
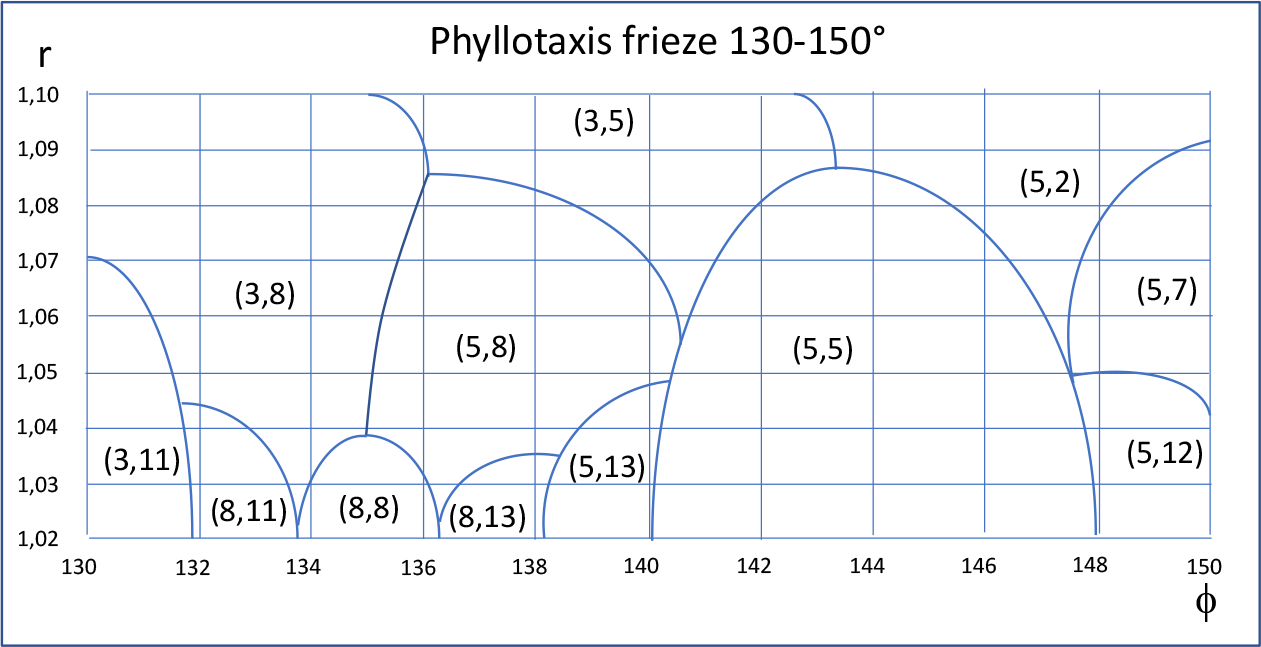
Phyllotaxis frieze delimiting the different phyllotaxis regions for divergence angles between 130° and 150°.

**Table 2.**
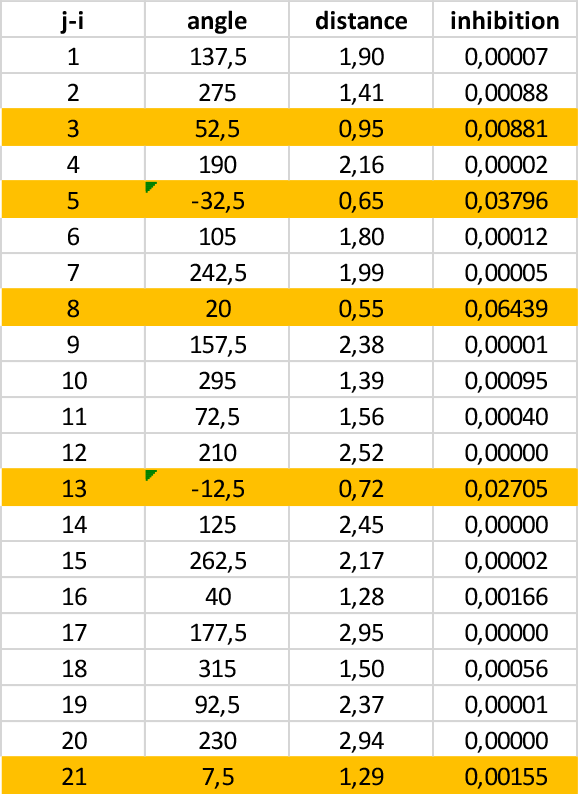
angular differences (between 0° and 360°) between a new primordium (i) and the already existing primordia (j) for a divergence angle of 137.5°; distances between these primordia for a plastochron ratio of 1.04, the radius R_0_ being taken as a unit of length; contribution to the inhibitory potential for *γ* = 0.2.

The phyllotaxis frieze is more readable than the van Iterson diagram, and offers a practical application: determining a phyllotaxis mode from the couple (ϕ, r) or vice versa a region (ϕ, r) from a phyllotaxis mode. In addition, it highlights the regions where a whorled or pseudo-whorled phyllotaxis such as (5,5) or (8,8) ensures a more dense distribution of primordia.

The model also makes it possible to calculate the value of the potential for any pair (ϕ, r). In the case γ = 0.2, only a few primordia participate actively in this potential.

**Figure 6.**
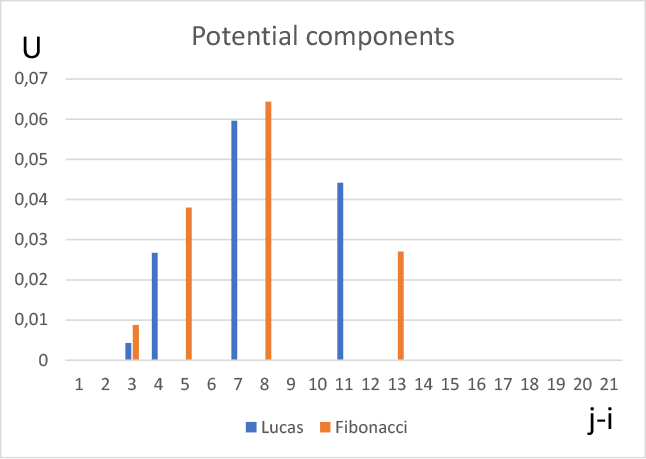
Potential components for the Fibonacci (137.5°) and Lucas (99.5°) angles for r = 1.04 and *γ* = 0.2. On the x-axis, the relative numbers of primordia (j-i; cf. table 2). For a divergence angle of 137.5°, it is obviously the closest primordia, 8, 5, 13 and 3 which produce the highest contribution to the total potential.

The total potential varies during meristem development depending on the divergence angle and the plastochron ratio.

**Figure 7.**
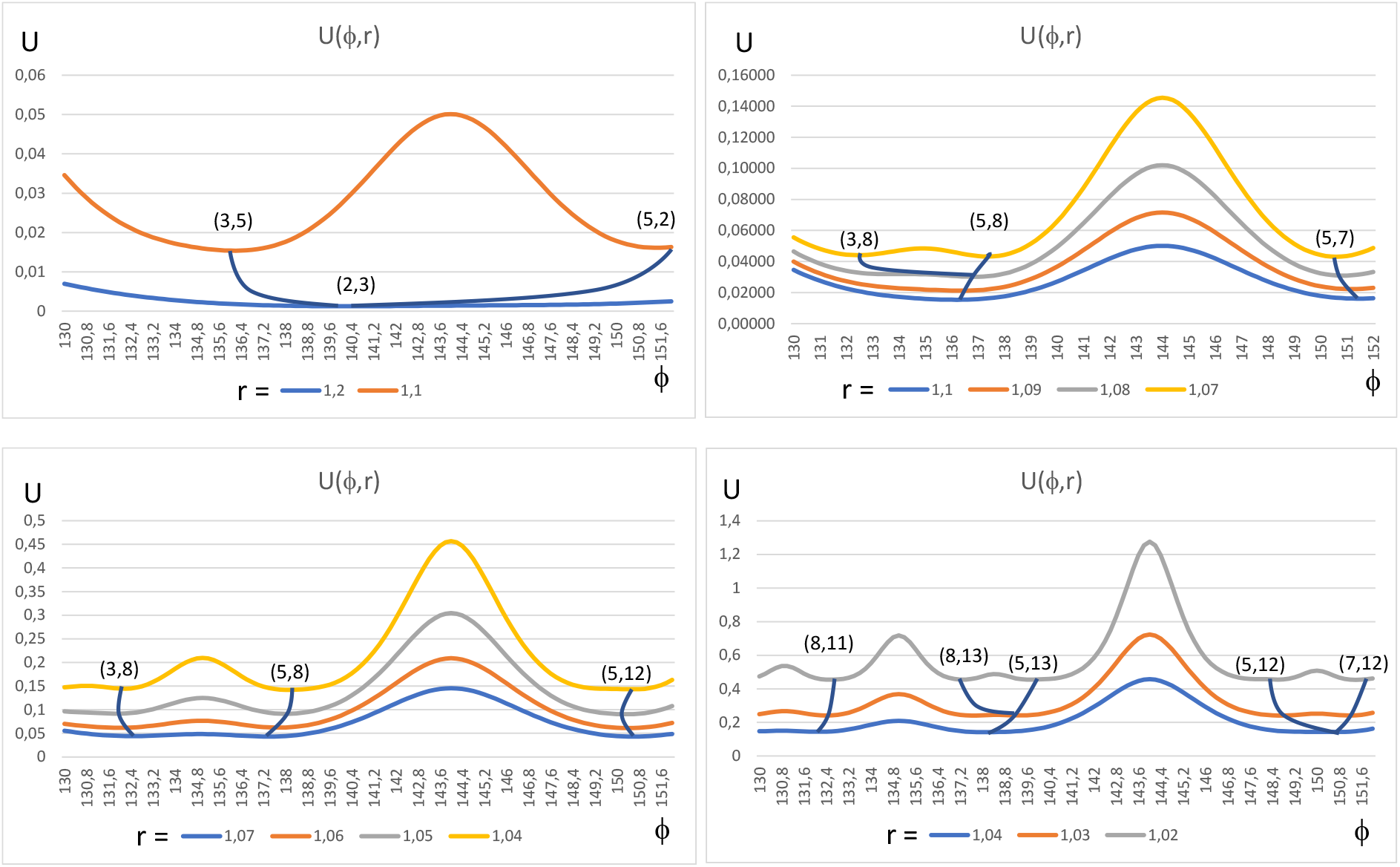
Values of the inhibition potential as a function of the divergence angle (ϕ) and the platochrone ratio (r) and trajectories in the phase space following the talwegs of potential.

Dynamics consistent with that of van Iterson diagram result from the movement of the system when the plastochron ratio decreases: primordia are initiated along the talwegs of potential, where the inhibition potential is minimum for a given plastochron ratio. We find the same bifurcations as van Iterson, the number of potential minima increasing with the decrease in plastochron ratios. If the trajectory in the direction of (8,13) corresponding to Fibonacci phyllotaxies is largely the most frequent in real plants (including *Linum*), phyllotaxis (8,11) has also been observed in *Magnolia* (Zagórska-Marek, 1994).

The divergence angles towards which phyllotaxies tend for very low values of the plastochron ratio are the “noble angles” (Jean, 1994). These angles, characterized by their representation in the form of “continuous fractions”, optimize the distribution of primordia within meristems (Marzec and Kappraff, 1983). These are, in the range of plastochron ratios studied here: 132.18°; 137.51°; 148.77° and 151.13°. These values correspond to the limit of an infinite sequence whose terms are found in the phyllotaxis mode produced when the plastochron ratio decreases (number of primordia to make n turns) divided by the corresponding term of the Fibonacci sequence (number n of laps performed). For example, 12 + 7 = 19 divided by 3 + 5 = 8 gives 0.421 turn, or 151.58°, an approximation of our fourth noble angle.

## 5 The sequence of plastochron ratios

According to the models of Douady and Couder (1996b) or Smith and al. (2006), just after the initiation of the primordium i-1, when the meristem continues to grow, the potential at any point of the front decreases. When at a point of this circle the potential falls below a threshold, the primordium i can be initiated. This condition is written:

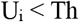

where Th (for threshold) is a constant. This condition is only realistic if the growth is stationary (r = constant). But if the plastochron ratio decreases, the potential at the front for a given divergence angle rises almost exponentially as the front is more and more crowded.

**Figure 8.**
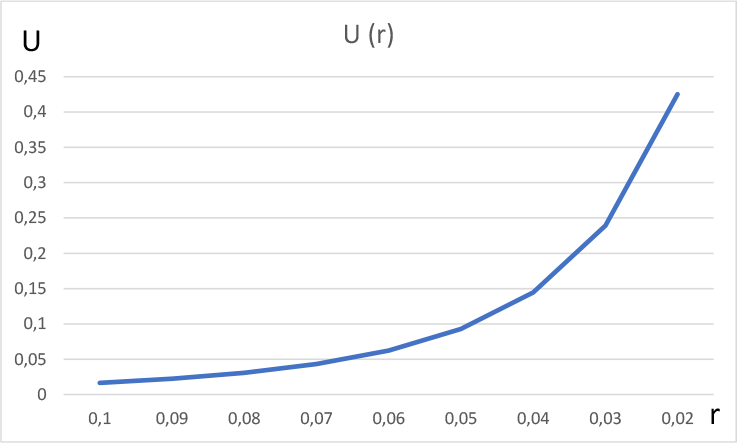
Value of the potential as a function of the plastochron ratio for a divergence angle of 137.5° and γ = 0.2 from equations (1) and (2).

We follow the hypothesis suggested by *Linum*’s modeling according to which a new primordium can be initiated when the difference between the potential U_i_ and the potential U_i-1_ become inferior to a given threshold, ΔUth.

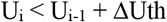

For a given divergence angle, knowing the potential as a function of the plastochron ratio, we can determine the number of primordia for different intervals of plastochron ratios: we divide the potential difference between the limits of each of these intervals by ΔUth.

The threshold ΔUth was determined so as to reproduce as accurately as possible the Williams data for *Linum*. For the first primordia (1-6), the sequence of plastochron ratios cannot be determined in the same way because the divergence angles are far from 137.5°.

The development of a meristem usually goes through three stages. First, the angular arrangement of the first primordia, to which the initial conditions of the model correspond. Then, the decreasing sequence of plastochron ratios, because any growth phenomenon is naturally limited. Finally, after the initiation of a large number of primordia, the plastochron ratios tend to stabilize (Figure 9). This is the “steady state” phase (Galvan-Ampudia, 2020).

**Figure 9.**
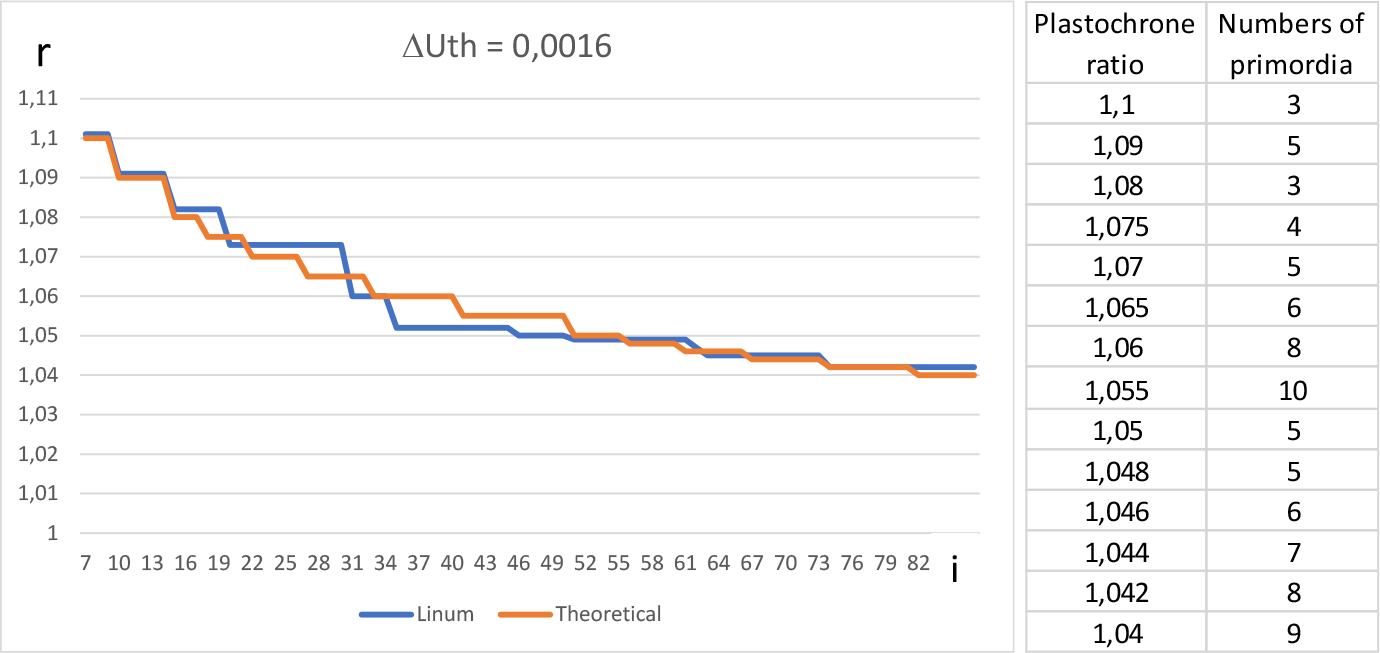
Sequence of plastochron ratios observed in flax (Williams, 1975) and according to the model for a divergence angle of 137.5° and a potential difference ΔUth = 0.0016.

## 6 Discussion

The van Iterson diagram traces the trajectories which lead to the different phyllotaxic modes. We based this diagram on a principle of minimizing the inhibitory potential under the constraint of decreasing plastochron ratios. This gives a totally different landscape of phyllotaxis morphogenesis from that resulting from the stacking of hard disks in a cylinder. According to Godin, Golé and Douady (2020), phyllotaxis is the consequence of a “geometric channeling” during plant development: spiral phyllotaxis would only be due to purely geometric constraints imposed by the competition of organs for space within the meristem. According to our approach, it is an emerging property of the physicochemical process implemented by the primordium initiation automaton. The mutations that distinguish one species from the others modify the parameterization of this automaton, that is to say the sequence of plastochron ratios. This is what allows the great diversity of phyllotaxies.

If the van Iterson diagram traces the trajectories which lead to the different phyllotaxic modes, it gives no indication of the speed of real plants on these trajectories. By showing that the plastochrone ratio sequence measured by Williams on flax respects the values of the inhibition potential as a function of the plastochrone ratios, we have determined that it give a good approximation to the dynamics leading to Fibonacci phyllotaxies.

The divergence angles that are selected by minimizing the inhibitory potential optimize the distribution of primordia in the meristem. Strauss and al. (2020) found that the same angles maximize light captured by a plant’s foliage by minimizing the shade of a leaf on those below. However, the golden angle does not perform better than other noble angles, which leaves open the question of its preponderance among stem and floral meristems.

The form of the inhibitory potential used here has no physicochemical basis. Douady and Couder (1996a) have shown that different potentials, provided they are realistic, lead to qualitatively similar results. The hypotheses of our model are only credible insofar their observable consequences conform to botanical reality. Chemical scientists themselves are sometimes forced into such empiricism, for example when they use the Lennard-Jones potential, which describes the interactions between two atoms of a monatomic gas (noble gas). At short distance, these prevent the mutual interpenetration of the electron clouds of the two atoms: the formula of this potential is purely empirical.

The fit between our model and the data measured by Williams on the flax meristems is outstanding. Nevertheless, many models of real plants must be carried out to support any methodological approach. That based on plastochron ratios had already been implemented by Kitazawa and Fujimoto (2015) who modeled the floral meristem of *Silene*. We have given it basis through the relationship between inhibition threshold and plastochron ratios. Other species modelling are in progress, based on a good communication between botanists and physicists.

## Acknowledgments

We gratefully acknowledge Solange Blaise (Société Botanique de France) for her useful discussions about botanical issues of phyllotaxis.

